# Distinct sources of deterministic and stochastic components of action timing decisions in rodent frontal cortex

**DOI:** 10.1101/088963

**Authors:** Masayoshi Murakami, Hanan Shteingart, Yonatan Loewenstein, Zachary F. Mainen

## Abstract

The selection and timing of actions are subject to determinate influences such as sensory cues and internal state as well as to effectively stochastic variability. Although stochastic choice mechanisms are assumed by many theoretical models, their origin and mechanisms remain poorly understood. Here we investigated this issue by studying how neural circuits in the frontal cortex determine action timing in rats performing a waiting task. Electrophysiological recordings from two regions necessary for this behavior, medial prefrontal cortex (mPFC) and secondary motor cortex (M2), revealed an unexpected functional dissociation. Both areas encoded deterministic biases in action timing, but only M2 neurons reflected stochastic trial-by-trial fluctuations. This differential coding was reflected in distinct timescales of neural dynamics in the two frontal cortical areas. These results suggest a two-stage model in which stochastic components of action timing decisions are injected by circuits downstream of those carrying deterministic bias signals.

## INTRODUCTION

Decisions are influenced by many factors including sensory information relevant to inferring actions and possible outcomes, the subjective values of those outcomes and the internal state of the chooser. Such factors allow, to some degree, choices to be predicted and can be considered **deterministic** factors. However, typically the totality of all available deterministic factors only accounts for a tendency or probability of choice (Sugrue et al., 2005). In order to enact an actual choice, additional covert mechanisms appear to determine, according to these probabilities, what action is taken and when. These processes contribute an effectively **stochastic** component to choice behavior. Beyond simply resolving indeterminacy, stochastic factors can also have adaptive value, such as helping to fairly explore the landscape of possible actions to aid adaptive learning (Fee and Goldberg, 2011) or adding unpredictability to behavior in competitive interactions (Barraclough et al., 2004; Dorris and Glimcher, 2004; Tervo et al., 2014). Biased stochastic choice models are ubiquitous in computational theory, including accumulator or diffusion to bound models (Gold and Shadlen, 2007; Ratcliff and Rouder, 1998) and reinforcement learning models (Dayan and Abott, 2001; Sutton and Barto, 1998). However, we still do not know how such processes are actually supported by neural circuits. In fact apparently stochastic components might reflect in part the deterministic influences of unobserved factors such as reward-dependent learning (Mendonca et al., 2012).

The deterministic and stochastic components of a decision process are relevant not only to the selection of actions, but also to the determination of *when* to act. An interesting example of a “when” decision occurs when an agent is waiting for a delayed and unpredictable event. Here a decision about how long to wait or when to give up waiting takes place. Such waiting decisions can be biased by the value associated with the delayed event, the history of past event times, the subjects’ temporal discounting factor and the internal state of the subject. Such deterministic factors bias the agent toward shorter or longer waiting times. In addition, the waiting decision may be also governed by effectively stochastic components that can result in substantial variability across multiple trials, even in a statistically stable environment (Murakami et al., 2014). Such variability can be captured by, for example, stochastic diffusion to bound models (Murakami et al., 2014; Schurger et al., 2012).

Murakami et al. (2014) found that trial-to-trial waiting times are robustly represented in the neural activity of the rat M2. The mPFC, an area reciprocally connected with M2 (Hoover and Vertes, 2007), is also implicated in waiting behavior. Lesions or inactivation of mPFC has repeatedly been shown to impair the ability of subjects to withhold responses for a delayed reward (Muir et al., 1996; Narayanan et al., 2006; Risterucci et al., 2003). Neural recordings have also found correlates of waiting behavior in the mPFC (Donnelly et al., 2015; Narayanan and Laubach, 2006). Thus, both M2 and mPFC appear to be highly relevant to waiting decisions, but it is unclear how neural populations in these two areas might be differentially involved.

In this study, we sought to understand how deterministic and stochastic components of decision processes are represented within neural populations in the M2 and mPFC. To do so, we used a task in which rats are required to wait for a randomly delayed tone to obtain a large reward, but also allowed to give up at any time to obtain a smaller reward (Murakami et al., 2014). Here, we found that waiting times could be well described by a two factor model. A deterministic component of waiting could be deduced from the recent history of waiting and rewards. A stochastic component could be defined as the residual trial-to-trial variability remaining after the deterministic bias was subtracted out. The deterministic signal had a much slower rate of change than the stochastic signal, allowing us to decouple them and search for their respective neural basis.

Inactivation experiments showed that both M2 and mPFC contribute to waiting times. Using multiple single-unit recordings, we found that stochastic waiting time variability was robustly encoded in M2, but not mPFC. In contrast, neural correlates of the deterministic history-dependent waiting time bias were found in both areas. But the deterministic signals had distinct temporal dynamics in M2 and mPFC: while these signals appeared transiently in individual M2 neurons, they were much more sustained in mPFC neurons, spanning throughout both the trial and inter-trial interval. The temporally distinct functions of the two areas were associated with distinct timescales of neural dynamics, demonstrated using firing rate autocorrelation analysis. Our results support the notion of a two-stage decision process. mPFC appears to be part of a network that maintains a slowly-varying, deterministic choice bias while M2 is part of a different system that translates this deterministic bias signal into actual choice signals while introducing an effectively stochastic trial-to-trial variability.

## RESULTS

### Behavior in a waiting task

Rats were trained to wait while holding their snout in a waiting port (Figure 1A) (Murakami et al., 2014). After a short delay (0.4 s), a first tone (T1) was played, after which the rat could garner a small amount of water reward by moving to an adjacent reward port. However if the rat chose to wait until a randomly-delayed second tone (T2, exponential distribution), it could garner a larger reward at the same reward port. Trials could be classified into three types according to rats’ waiting times in relation to presentation of tones: “short-poke” trials in which a rat failed to wait until T1 (7.6 ± 4.6%, mean ± s.d., *N* = 33 rats), “impatient” trials in which the rat waited past T1 but not T2 (59.0 ± 4.4%) and “patient” trials in which the rat waited beyond T2 (33.4 ± 1.3%).

**Figure 1.**
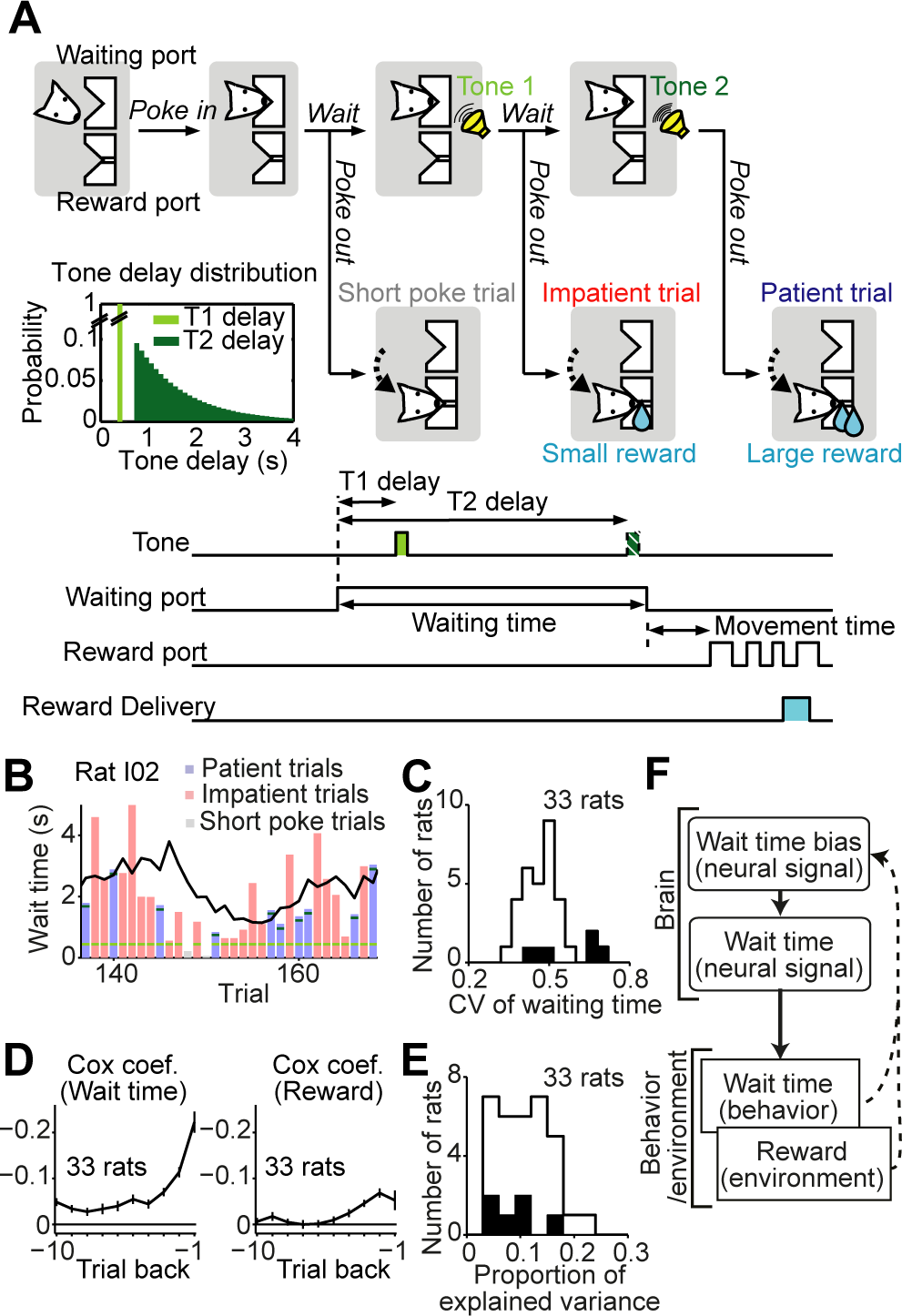
Two-Stage Decision Model for a Waiting Task. (A) Schematic of trial events in the waiting task (top). In each trial, after waiting for a certain period at the waiting port, a rat received tone(s), moved to the reward port and received a water reward, the size of which depended on the number of the tones presented. Inset, probability distributions of the delays to tone 1 (T1, light green) and tone 2 (T2, dark green). Bottom. Timeline of the trial events and the definition of the behavior parameters. The upper position of the lines for tone, waiting port, reward port and reward delivery indicates the presentation of tone(s), staying at the waiting port, staying at the reward port and reward delivery, respectively. The light green rectangle and dark green rectangle indicate tone 1 and tone 2, respectively. Tone 2 is represented by a hatched rectangle to indicate it was not played in the impatient trials. (B) Snapshot of the waiting behavior. Each bar indicates the waiting time of each trial. The black line indicates the waiting time bias estimated with the Cox regression model. Gray bars indicate short poke trials, pink bars impatient trials and blue bars patient trials. Light green ticks and dark green ticks represent tone 1 and tone 2, respectively. (C) A histogram of the coefficient of variation (CV) of waiting time across trials for 33 rats. Filled bars indicate rats used for neural data analyses. (D) Cox regression analysis of the waiting time. Shown are Cox regression coefficients for the waiting time (left) and reward (right) of previous trials (1 to 10 trial back). Average across rats (*N* = 33 rats). Error bars indicate the standard error. Note that negative Cox coefficients mean a negative effect of the predictors on the probability of giving up waiting, which eventually means a positive effect on the waiting time. (E) A histogram of proportions of variance explained (Schemper’s V)(Schemper and Henderson, 2000) with the Cox regression model for 33 rats. Filled bars indicate rats used for neural data analyses. (F) A schematic diagram of the waiting time decisions.

Rats’ waiting times showed substantial trial-to-trial variability (Coefficient of Variation, CV: 0.48 ± 0.08 s, Figures 1B and 1C). We first determined what fraction of this variability could be predicted by trial history. To quantify how trial history affects the waiting time of individual trials, we performed a regression analysis using past trial waiting time and reward history as regressors. We used a type of regression analysis, Cox regression analysis, suitable for time-to-event data. Cox regression accounts for interrupted trials, allowing both patient and impatient trials to be included in the analysis. Cox regression revealed significantly negative coefficients for both waiting time and rewards of previous trials (Figure 1D). That is, longer waiting times and larger rewards garnered in previous trials decreased the probability of giving up waiting, lengthening waiting times. The Cox regression model explained on average 11.1 ± 5.0% (*N* = 33 rats) of variability in waiting behavior (Figure 1E). Although modest, contributions of both waiting time and prior rewards were significant in most of the rats (*P* < 0.05, for any of the coefficients from 1 to 3 trials back, correcting for multiple comparisons, 31/33 and 22/33 rats respectively). Note that although additional slowly-varying factors such as thirst state are also likely to contribute to slow trial-to-trial variations in waiting, these factors would introduce waiting time trial-to-trial autocorrelation (Figure S1) and therefore should be effectively captured by the coefficient for past trial waiting time. These results could be described by a two-stage decision model in which at the beginning of each trial, rats have a deterministic bias in waiting time, but the actual waiting time is specified by a process that adds stochastic trial-to-trial variability. The resulting waiting behavior and reward outcome of the current trial would in turn update the deterministic waiting time bias in the following trial (Figure 1F).

### Pharmacological inactivation revealed necessity of both mPFC and M2 in a waiting task

To test the causal influence of area M2 and mPFC in the waiting task, we examined behavioral performance after inactivating these areas in alternating sessions in which either vehicle or the GABA-A agonist muscimol were injected bilaterally. Inactivation of M2 caused a dramatic impairment in performance. The direction of the effect on waiting behavior was not consistent across rats (waiting time: *P* = 0.28 repeated measures ANOVA or rmANOVA; fraction of patient trials: *P* = 0.083, rmANOVA followed by t-tests, Figure 2A and 2B). However, at an individual animal level, all the rats showed a significant change in the waiting time distribution (*P* < 0.05, Kolmogorov-Smirnov test) and most of the rats showed either increase or decrease in the waiting time (*P* < 0.05, one-way ANOVA followed by two-sample t-tests, Figure 2A). Inactivation of M2 also caused more general impairments in performance: the time to move to the reward port increased (*P* < 0.05, rmANOVA followed by t-tests, Figure 2C) and the fraction of reward-port-visit trials decreased (*P* < 0.05, rmANOVA followed by t-tests, Figure 2D). Although the changes in various aspects of behavior prevent us from specifying a single primary effect of M2 inactivation, the result showed the necessity of M2 in performing this task.

**Figure 2.**
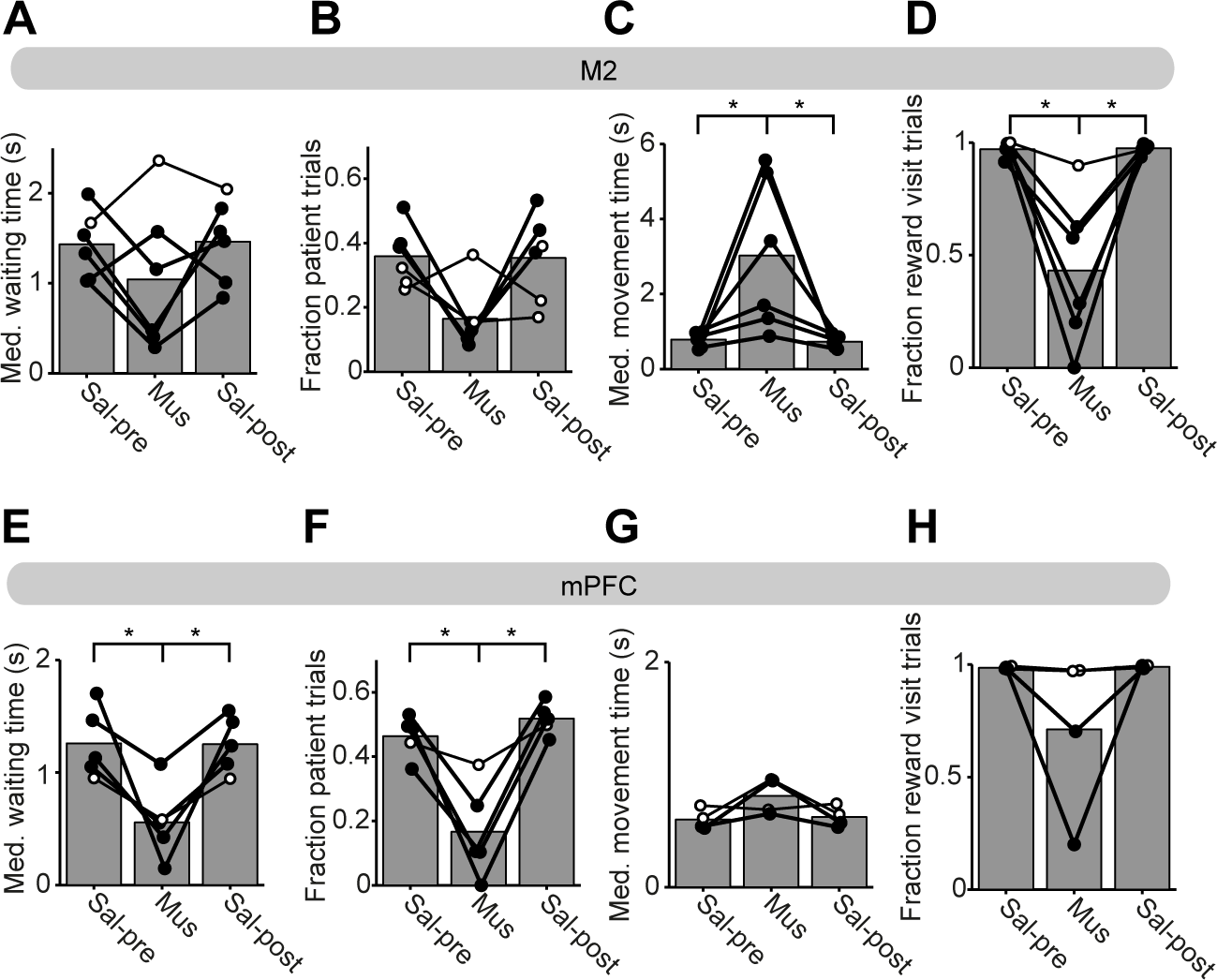
M2 and mPFC are Necessary for Performing the Waiting Task. (A) The bars represent across-rat averages of the median waiting time from the control session before the muscimol session (Sal-pre, left), the M2 inactivation session (Mus, middle) and the control session after the muscimol session (Sal-post, right). Circles connected by a line indicate an individual rat. Filled circles indicate rats showing a significant difference between inactivation session and control sessions. Open circles indicate a rat without significant difference. (B) The same as (A), but for the fraction of patient trials. (C) The same as (A) but for the median movement time. Asterisks indicate significant difference between inactivation session and control sessions (*P* < 0.05, repeated measures ANOVA, followed by paired t-tests). (D) The same as (A) but for the fraction of reward visit trials. (E-H) The same as (A-D) but for mPFC inactivation. Asterisks indicate significant difference between sessions (*P* < 0.05, rmANOVA followed by paired t-tests).

Bilateral inactivation of mPFC also caused severe impairments in performance of the waiting task, but with different spectra of behavioral changes compared to the M2 inactivation. mPFC inactivation consistently impaired waiting performance in all rats, as indexed by a decrease in waiting time (*P* < 0.05, rmANOVA followed by t-tests, Figure 2E) and in the fraction of patient trials (*P* < 0.05, rmANOVA followed by t-tests, Figure 2F). This impairment in waiting behavior was independent of motor effects: the movement time and fraction of reward-port-visit trials did not change significantly between inactivation and control sessions (movement time: *P* = 0.084, rmANOVA, fraction of reward-port-visit: *P* = 0.18, rmANOVA, Figure 2G and 2H). In summary, inactivation experiments suggest that both M2 and mPFC are crucial for performing the waiting task. Despite the overall similarity in the effect of inactivation of both areas, differences in the pattern of behavioral effects suggest differential contributions of these areas to the waiting task. We next explored this idea further using electrophysiological experiments.

### M2, but not mPFC, encodes actual waiting times

To search for neural signals involved in waiting time decisions, we performed chronic multiple single-unit recordings from M2 and mPFC using tetrode arrays. The data set consisted of 306 neurons in M2 (average of 5.0 ± 5.4 neurons/session, 61 sessions over 5 rats) and 126 neurons in mPFC (average of 4.8 ± 3.8 neurons/session, 26 sessions over 3 rats). We first looked for neural signals involved in the final “implementation” stage of waiting time decisions, i.e. neural signals correlated with the actual waiting time. As shown previously, a significant fraction of M2 neurons exhibited waiting-time-correlated activity (Murakami et al., 2014). Some neurons showed ramping activity whose threshold-crossing time was predictive of waiting times (ramp-to-threshold type, 11%, *P* < 0.001, one-sided permutation test with trial shuffling, Figure 3A and 3C). Other neurons exhibited transiently waiting time correlated activity (transient correlation type, 16%, *P* < 0.001, one-sided permutation test, Figure 3D and 3F). In striking contrast, the activities of mPFC neurons showed neither type of waiting time correlation: Ramp-to-threshold analysis showed zero neurons predictive of the waiting time (0%, *P* = 1, one-sided permutation test, Figure 3B and 3C). Even when looser criteria to define ramp-to-threshold type neurons were used (see Experimental Procedures), we only found a single additional neuron (1/126 neurons). Similarly, the transient correlation analysis found few waiting-time predictive neurons (5.6%, *P* = 0.69 with one-sided permutation test, Figure 3E and 3F), not more than expected by chance.

**Figure 3.**
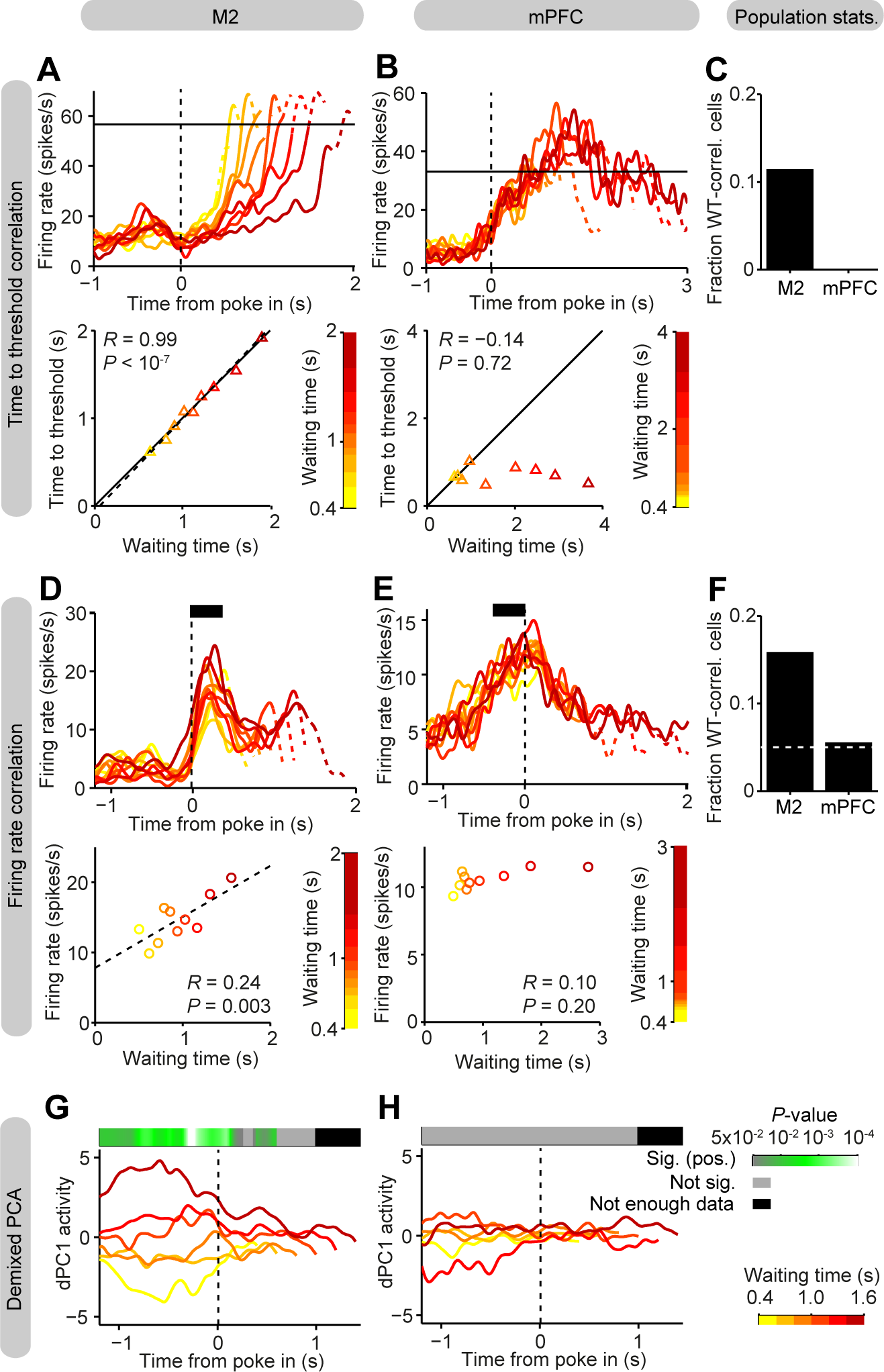
M2, but not mPFC, Encodes the Waiting Time. (A) Top. Spike density functions (SDF) of an example ramp-to-threshold type M2 neuron for different waiting time trials, aligned to poke-in. Impatient trials are grouped according to the waiting time, indicated by the color scale at the bottom. Bottom. Time to cross an example threshold level, indicated by a horizontal line at the top panel, as a function of mean waiting time. The dashed line indicates the regression line. (B) SDF of an example mPFC neuron (top) and time to cross a threshold level as a function of mean waiting time (bottom). Conventions are the same as in (A). (C) Fraction of ramp-to-threshold type neurons in M2 and mPFC. (D) Top. SDF of an example transient correlation type M2 neuron. Conventions are the same as in (A). Bottom. Firing rate at an example 0.4 s time window, indicated by a filled bar at the top panel, as a function of mean waiting time. The dashed line indicates the regression line. (E) SDF of an example mPFC neuron (top) and firing rate at a 0.4 s time window as a function of mean waiting time (bottom). Conventions are the same as in (D). (F) Fraction of transient correlation type neurons in mPFC and M2. The dashed line indicates the chance level of 0.05. (G) Activity (score) of the first demixed principal component (dPC1) of M2 neurons, aligned to poke-in. Shown are dPC1 activities from impatient trials with different waiting times, indicated by the color scale on the right. The top bar indicates *P*-value of Pearson’s correlation coefficient calculated at each time point. The color scale of the *P*-value is indicated on the right. Green for significant positive correlation, gray for not significant, black for not enough data. (H) dPC1 activity of mPFC neurons. Conventions are the same as in (G).

Although we did not find waiting time correlated signals in mPFC in individual neuron analyses, such signals might be hidden in population activity. In order to search for such signals, we performed demixed principal component analysis (dPCA) (Kobak et al., 2016). This analysis, in essence, finds linear combinations of neurons whose population activity varies along parameters of interest (demixed principal components, dPCs), which in this case was only the actual waiting time. Using dPCA, consistent with the single unit analysis, we found waiting time correlated signals in M2 (Figure 3G). The first component, dPC1, showed strong correlation with waiting times before the waiting period, while the second and third (dPC2 and dPC3) showed correlations during waiting period (Figure S2A). On the other hand, and again consistent with the single neuron analyses, we failed to find waiting time correlated signals in mPFC population activity (Figure 3H and S2B). In mPFC, dPC activities from different waiting time trials largely overlapped with each other and were not consistent across time. Controlling the difference in a number of neurons used in the dPCA for M2 and mPFC did not explain the strength of wait time coding (Figure S2C and S2D). Taken together, both the single neuron analyses and population analyses suggest that the actual waiting time is robustly encoded in the M2 neurons, but not in the mPFC neurons, at least not in linearly decodable form.

### Both M2 and mPFC encode history-dependent, deterministic waiting time biases

Next, we looked for neural signals encoding deterministic waiting time biases. To do so, we examined the correlation between neural activity and deterministic waiting time bias estimated using the Cox regression (Figure 1B and 1D). Figure 4A shows an example M2 neuron that was strongly excited just after poke-in. The amount of activation was significantly correlated with the waiting time bias (*R* = 0.41, *P* = 0.0001, Figure 4A). Similarly to the M2 neurons, mPFC neurons also showed waiting time bias-correlated activity. In mPFC, we also found a neuron whose activity was correlated with the waiting time bias (Figure 4B). This example mPFC neuron did not show any transient modulation of neural activity around poke-in period but had a level of ongoing activity that differed across trials with different waiting time bias. The neuron’s firing rate was elevated in trials with a long waiting time bias compared to trials with a short waiting time bias. The firing rate was strongly correlated with the waiting time bias (*R* = 0.37, *P* = 0.0002). As a population, 11% of M2 neurons and 16% of mPFC neurons showed waiting time bias-correlated activity (*P* < 0.001 for M2 and mPFC, one-sided permutation test with trial-shuffling, *P* < 0.05 for M2, *P* < 0.01 for mPFC, one-sided permutation test with session-shuffling, Figure 4C). We also performed dPCA, now targeting the deterministic waiting time bias. Consistent with single neuron analyses, dPCA also revealed strong waiting time bias signals in both populations of M2 and mPFC (Figure 4D and 4E). These results indicate that whereas only M2 neurons encode actual waiting times, both M2 and mPFC neurons encode deterministic waiting time bias signals.

**Figure 4.**
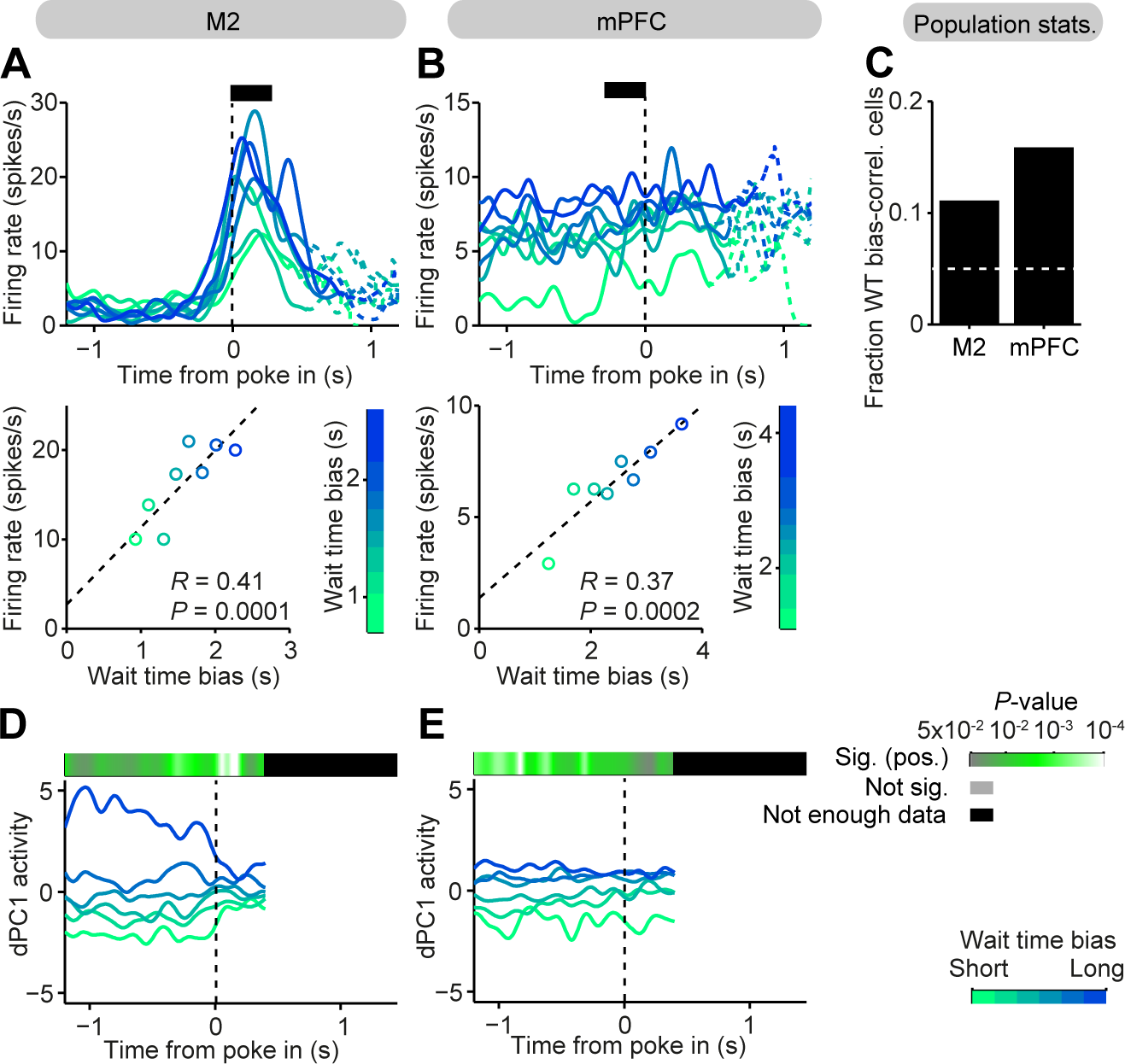
Both M2 and mPFC Encode the Waiting Time Bias. (A) Top. Spike density functions (SDF) of an example M2 neuron for trials with different waiting time bias, aligned to poke-in. Impatient trials are grouped according to the waiting time bias, indicated by the color scale at the bottom. Waiting time bias is calculated with Cox regression including 1 – 5 trial back trial histories. Bottom. Average firing rates for each group of trials at an example 0.4 s time window indicated by a filled bar in the top panel, as a function of mean waiting time bias. The dashed line indicates the regression line (*R* = 0.41, *P* = 0.0001). (B) SDFs of an example mPFC neuron (top) and average firing rates at a 0.4 s time window indicated by a bar in the top panel as a function of mean waiting time bias (bottom, *R* = 0.37, *P* = 0.0002). Conventions are the same as in (A). (C) Fraction of waiting time bias correlated neurons in M2 and mPFC. Dashed line indicates the chance level of 0.05. (D) Activity (score) of the first demixed principal component (dPC1) of M2 neurons, targeting the waiting time bias signals, aligned to poke-in. Shown are dPC1 activities from impatient trials with different waiting time biases, indicated by the color scale on the right. The top bar indicates *P*-value of Pearson’s correlation coefficient calculated at each time point. The color scale of the *P*-value is indicated to the right. Green for significant positive correlation, gray for not significant, black for not enough data. (E) dPC1 activity of mPFC neurons, targeting the waiting time bias. Conventions are the same as in (D).

Deterministic waiting time biases are correlated with actual waiting times. Thus, M2 deterministic waiting time bias signals might be simply due to the existence of M2 neurons with actual waiting time signals. Alternatively, the neurons encoding the deterministic bias might form a separate population from neurons encoding actual waiting time. To test this, first, we decomposed actual waiting times into a trial-history-dependent, deterministic “bias” component and stochastic “residual” component. Then, we performed a multiple linear regression analysis in order to estimate the extent to which neural activity is influenced by the deterministic bias and stochastic residual factors. We found that 16% of M2 neurons carried the bias signal and 16% carried the residual signal (Figure 5A, see Experimental Procedures). We then tested whether these two populations form separate populations or overlapping populations. We found that regression coefficients for the bias and that for the residual component were positively correlated (*R* = 0.37, *P* < 10^−10^, Figure 5B). This suggests that the M2 neurons tend to carry the bias signal and the residual waiting time signal together. A parsimonious explanation for this result is that M2 neurons carry the actual waiting time, that is, the stochastic trial-to-trial variability added on top of the deterministic waiting time bias. However, there were also subpopulations of M2 neurons that encoded strongly the deterministic bias, but not the stochastic residual waiting time, and vice versa. These subpopulations appear to form a continuum rather than discrete populations.

**Figure 5.**
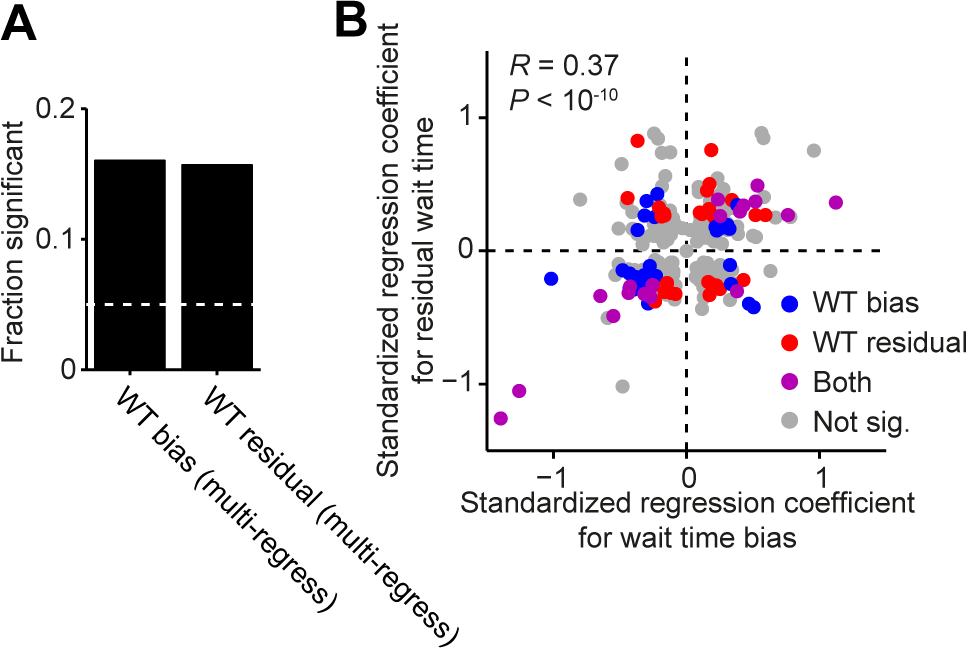
Distribution of the Waiting Time Bias and Residual Waiting Time Signals in the M2 Population. (A) Fraction of waiting time bias neurons and residual waiting time neurons detected with a multiple linear regression analysis. The dashed line indicates a chance level of 0.05. (B) A scatter plot representing a standardized regression coefficient for the waiting time bias (x-axis) and that for the residual waiting time (y-axis) for each neuron. The regression coefficient was taken at the most significant bin for each variable. Red neurons indicate waiting time bias neurons, blue neurons residual waiting time neurons, purple neurons significant for both and gray neurons significant for neither.

### M2 and mPFC differ in timescales of waiting time coding

If M2 and mPFC contribute to different components of choice behavior, then given that these components unfold over different timescales, those contributions might be visible in the dynamics of their respective neural activity. To test this idea, we measured spike-count auto-correlation, previously used to measure intrinsic timescales of primate cortical areas along the cortical hierarchy (Murray et al., 2014). We indeed found that the auto-correlation function decayed more slowly in mPFC neurons compared to M2 neurons (Figure 6A, τ_M2_ = 373 ms, τ_mPFC_ = 514 ms, *P* < 0.001). When auto-correlation analysis was extended to the next trial, which normally occurred more than 10 seconds apart, mPFC neurons showed higher auto-correlation compared to M2 (Figure 6B, correlation coefficient between trial N and N+1, M2: 0.086; mPFC: 0.13, *P* = 0.002). These results indicate that mPFC has a slower timescale of neural dynamics, suitable to maintain the waiting time bias signal throughout a trial and across multiple trials. We also quantified the timescale, or more precisely “trial-scale”, on which neural activity co-fluctuates with waiting behavior. To do so, we measured coherency between waiting times and neural activity (see Experimental Procedures). For the M2 population, coherency between waiting times and neural activity was significant (compared to session-shuffled data) across a wide range of timescales, ranging from 2 trials to > 60 trials (Figure 6C). On the other hand, for mPFC, coherency was significant only at slower timescales, > 7 trials (Figure 6D). These results corroborate the idea that M2 and mPFC differ in encoding two stages of waiting decisions: only M2 neurons show correlations with behavior on the timescale of rapid stochastic waiting time fluctuations.

**Figure 6.**
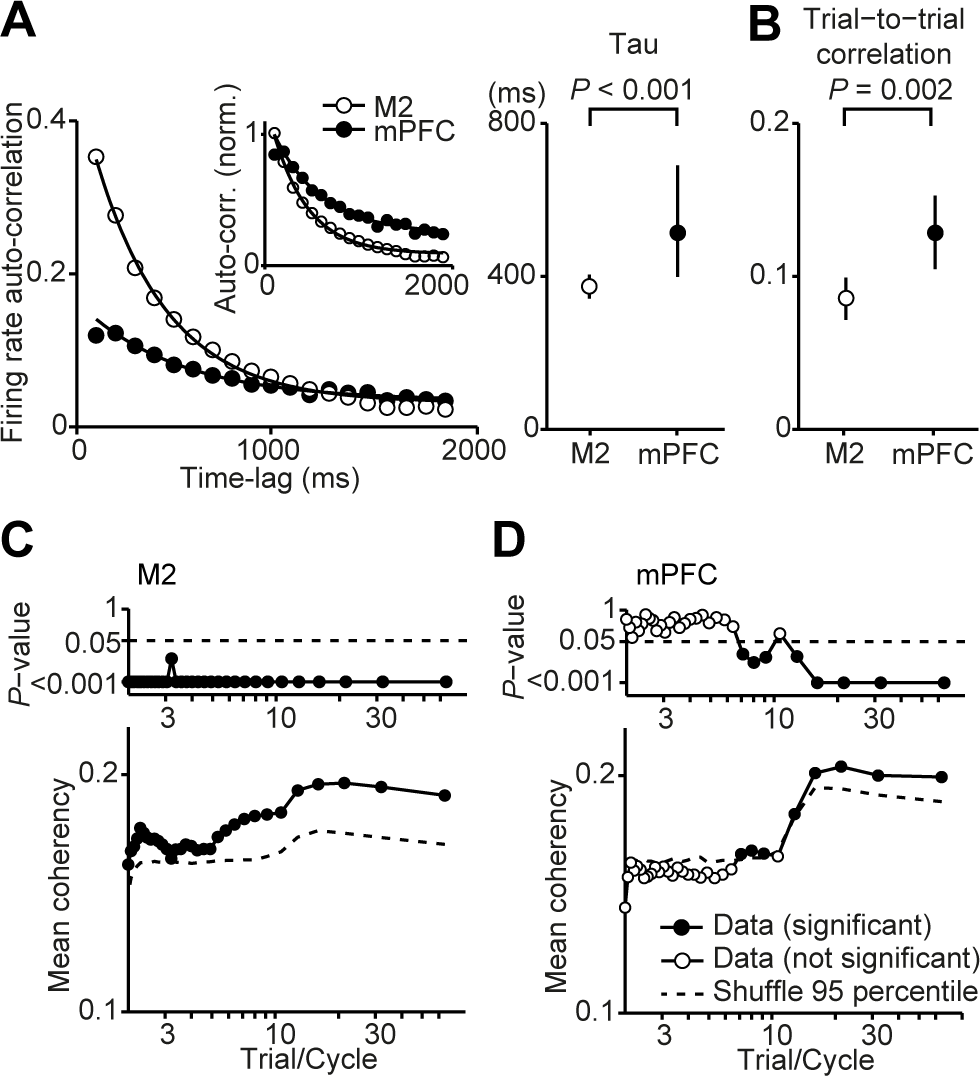
Timescale of mPFC and M2 Neural Dynamics. (A) Left. Spike-count auto-correlation functions for mPFC and M2 neurons. The auto-correlation analysis is performed at a 2 second window before the poke-in. Each line is an exponential fit for population of mPFC or M2 neurons. Shown in the inset is the auto-correlation function normalized to the value of exponential fit at time-lag = 100 ms. Right. Decay parameter, tau, of the exponential fit for mPFC and M2 neurons. The error bars indicate bootstrap 95% confidence intervals. *P*-value is calculated with a permutation test. (B) Spike-count auto-correlation between two consecutive trials for mPFC and M2 neurons. Spike-count is measured at 2 second window before the poke-in. The error bars indicate bootstrap 95% confidence intervals. *P*-value is calculated with a permutation test. (C) Bottom. Coherency between the waiting time and M2 firing rate across trials. Average coherency across time bins and across neurons (261 neurons, 1044 bins). The dashed line indicates 95 percentile of session shuffle data. Top. *P*-value of the mean coherency estimated from a permutation test (one-sided test). (D) Bottom. Coherency between the waiting time and mPFC firing rate across trials. Average coherency across time bins across neurons (97 neurons, 388 bins). Top. *P*-value of the mean coherency estimated from a permutation test (one-sided test).

### mPFC waiting time bias signal persists throughout a trial and an inter-trial interval

The spike-count auto-correlation analysis and coherence analysis suggests that M2 and mPFC contribute differentially to choice behavior in part due to different intrinsic dynamics. Consistent with this notion, the example neurons in Figure 4A and 4B showed different temporal dynamics: While the example M2 neuron showed a waiting time bias signal in a transient activity burst, the mPFC neuron carried the same signal in a sustained activity. To further this analysis, we extended the analysis after the end of waiting, when the rat visited the reward port and consumed the reward (Figure 7A and 7B). Over this longer time window, the difference between M2 and mPFC dynamics was even more striking. While the example M2 neuron was sensitive to waiting time bias only during a brief transient during the waiting period (Figure 7A and 7C), the mPFC neuron’s waiting time bias signal was maintained throughout the trial, from pre-waiting period to the inter-trial interval (Figure 7B and 7E). This tendency was true for the populations of M2 and mPFC neurons. Waiting time bias signals were transient in M2 neurons and decayed quickly in different trial phases (Figure 7D) whereas they were maintained throughout the trial phases in mPFC neurons (Figure 7F).

**Figure 7.**
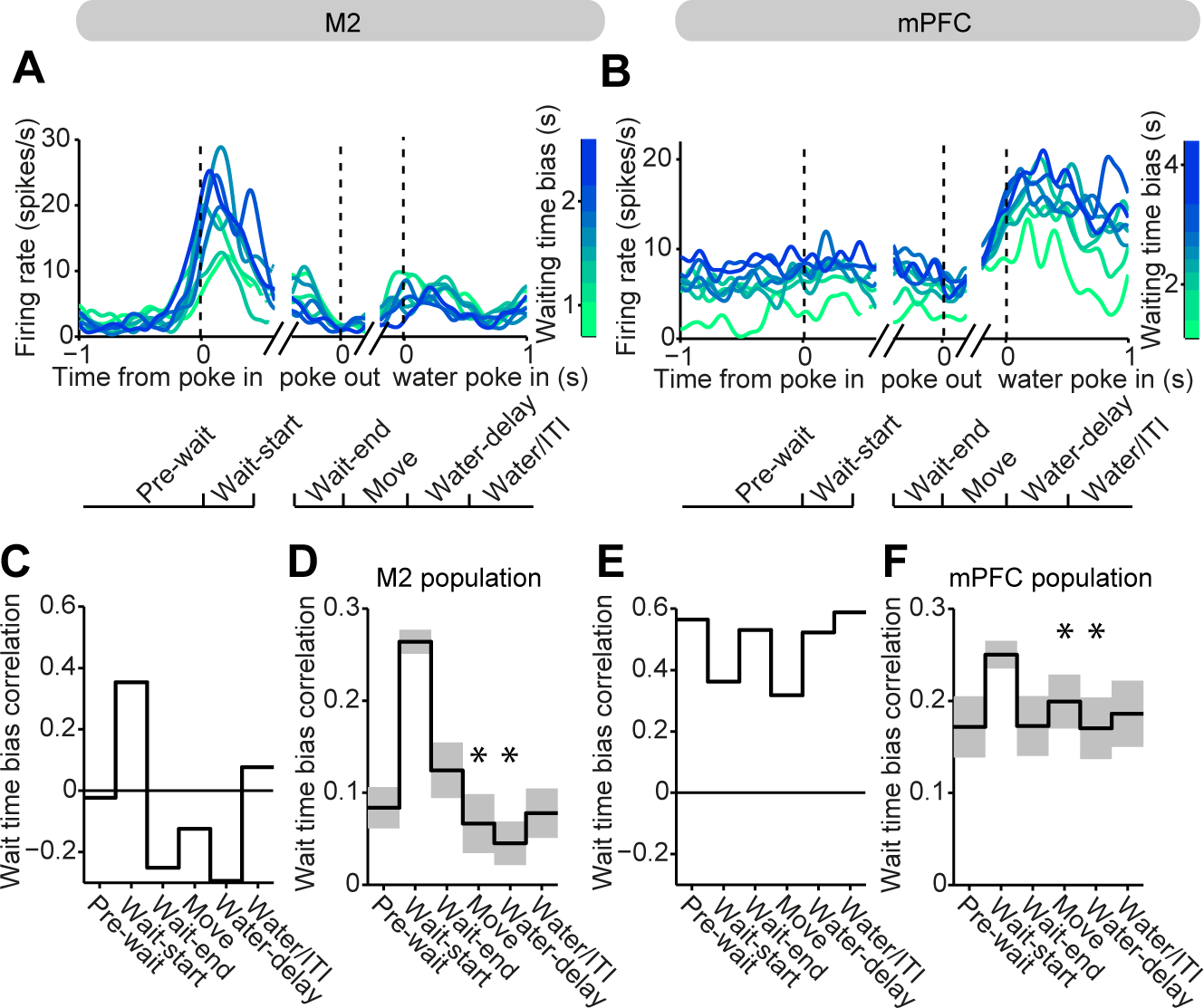
Different Temporal Dynamics of the Waiting Time Bias Signal in M2 and mPFC. (A) Spike density functions (SDF) of an example M2 neuron (the same neuron as in Figure 4A) for different waiting time bias trials, aligned to poke-in, poke-out and water-poke-in. Impatient trials are grouped according to the waiting time bias, indicated by the color scale. Shown at the bottom is the definition of trial phases. (B) SDFs of an example mPFC neuron (the same neuron as in Figure 4B). The conventions are the same as in (A). (C) Shown are Pearson’s correlation coefficients between the waiting time bias and the example M2 neuron’s firing rate at different trial phases. The same neuron as in (A). (D) Pearson’s correlation coefficients between the waiting time bias and the firing rate at different trial phases. Mean (black) and S.E.M (gray) across M2 neurons (*N* = 40 neurons). Asterisks indicate significant differences between M2 neurons vs. mPFC neurons at corresponding trial phases (*P* < 0.05, 2-sample t-test, corrected for 5 multiple comparisons for different phases excluding the phase used for the neuron selection). Neurons are selected for this analysis if it exhibited a significant correlation between the waiting time bias and the firing rate at the *wait-start* phase. Using different phases for neuron selection yielded similar results (Figure S3). (E) Shown are Pearson’s correlation coefficients between the waiting time bias and the example mPFC neuron’s firing rate at different trial phases. The same neuron as in (B). (F) Mean Pearson’s correlation coefficients between the waiting time bias and the firing rate at different trial phases for mPFC neurons (*N* = 25 neurons). The same conventions as in (D). Asterisks indicate significant differences against M2 neurons at corresponding phases (*P* < 0.05, 2-sample t-test, corrected for multiple comparisons).

## DISCUSSION

In this study, we investigated the neural basis of waiting decisions, identifying two separable components. First is the process of setting a deterministic bias, a tendency to wait shorter or longer, which we inferred from the previous trial history of rewards and waiting times. Second is the process of specifying the actual timing of actions, an effectively stochastic remaining component. The main conclusion from our study is that the deterministic and stochastic components are differentially encoded in neural population in the frontal cortex. mPFC neurons are part of a network that maintains deterministic bias signals while the M2 neurons are part of a network that translates this signal into specific waiting time signals, while injecting trial-to-trial stochasticity.

### Role of mPFC in waiting behavior

The mPFC is implicated in behavioral inhibition or control of impulsivity (Dalley et al., 2011), especially when a subject is required to withhold responses for a delayed reward (Muir et al., 1996; Narayanan et al., 2006; Risterucci et al., 2003). We verified the causal involvement of mPFC in the waiting task using reversible pharmacological inactivation by the GABA-A agonist muscimol. We showed that inactivation of mPFC, centered at the prelimbic cortex, impaired waiting behavior by decreasing average waiting times and the fraction of patient (successful) trials. These effects were not due to a general motor impairment, because movement to the reward port was not affected. Because the spectrum of behavioral changes produced by M2 inactivation was different to that of mPFC inactivation, spreading of muscimol dorsally into M2 does not explain the behavioral effect of the mPFC inactivation. Lateral spread of the drug is likely constrained by the cingulum or forceps minor of the corpus callosum according to our preliminary observations of dye injection. Because the mPFC, including the anterior cingulate cortex, prelimbic cortex and infralimbic cortex, is a dorsoventrally elongated structure, it is unlikely that the drug spread ventrally beyond mPFC. Further studies will be useful to dissect the function of different subregions within the mPFC.

We found that a significant fraction of mPFC neurons encodes a waiting time bias signal, that is, a tendency to wait shorter or longer depending on experience (rewards and achieved waiting times) in the immediately preceding trials. We refer to this signal as a “deterministic” bias signal because it is the predictable component. Because the deterministic bias signal was stably maintained in individual neurons even while the animal progressed through different sensorimotor experiences, such as nose-poke waiting, moving to the reward port and drinking water, it is very unlikely that this signal is trivially explained by sensorimotor variables (Cowen and McNaughton, 2007).

It is an open question what underlying variables this deterministic bias signal represents. The mPFC receives inputs from motor-related structures such as M2, as well as from valuation systems such as the amygdala, orbitofrontal cortex and midbrain dopaminergic systems (Van Eden et al., 1987; Hoover and Vertes, 2007). Thus it is suitable to represent action-outcome contingencies and build adaptive behavioral strategies based on task experience. Consistent with this view, mPFC is involved in trial-history-dependent strategy and must be actively suppressed for an animal to switch to history-independent stochastic strategy (Tervo et al., 2014). Alternatively, the waiting time bias signals might represent internal states relevant to this task, such as thirst or willingness to exert effort (de Araujo et al., 2003; Price, 1999; Walton et al., 2002). These internal states would likely also change depending on the trial history and in turn affect the waiting time. These possibilities are not mutually exclusive and both the precise task-related parameters and general internal states might influence the mPFC activity and be used in downstream areas to affect waiting behavior.

Neural correlates of waiting behavior in mPFC were reported previously, but the observed signals occurred too late to causally affect trial-to-trial waiting decisions on a given trial (Donnelly et al., 2015; Narayanan and Laubach, 2006). Those signals might thus reflect signals related to monitoring of waiting behavior that, together with trial outcome signals, would be useful to update future waiting behavior (Narayanan and Laubach, 2008). In contrast, the waiting time bias signals reported here occurred early in a trial, sometimes even before the initiation of waiting (Figure 4 and 7). Therefore, the observation is consistent with the idea that the deterministic waiting time bias signal is used to control the waiting behavior in the current trial. If waiting time bias signals were estimated from the behavioral data in previous studies, one might find neural signals correlated with the waiting time bias that appear early enough to influence behavior in the current trial.

Despite the fact that mPFC neurons could predict to some degree waiting time, they accounted for only a relatively small variance in the actual waiting time because they entirely lacked signals correlated with substantial trial-to-trial fluctuations. Thus, we conclude that a stochastic component of trial-to-trial waiting time is specified outside the mPFC. The absence of stochastic trial-to-trial action timing signal but the presence of slowly changing decision bias signal in the mPFC is reminiscent of mPFC neural activity recorded during a value-based two-alternative choice task (Sul et al., 2010). This study showed that in the mPFC trial-to-trial action planning signals are weak, but that neural activity is modulated by block-dependent reward probabilities. Together with the present results, this suggests that the mPFC is part of a network that maintains deterministic decision biases based on integrated information from slowly varying internal variables but is not directly involved in selecting impending actions.

### Role of M2 in waiting behavior

We found that M2, but not the mPFC, signaled the actual waiting time, which in this task included not only the slowly fluctuating, deterministic waiting time bias but also substantial trial-to-trial stochastic variability. Using a multiple-regression analysis to estimate the influence of both the slow deterministic bias component and fast stochastic component, we found that both components were present and positively correlated in M2 neuron firing rates. Because M2 neurons carry both components, we suggest that the M2 creates the final action timing signal by combining the deterministic and stochastic components. Despite this overall tendency of M2 population, some M2 neurons carried exclusively the deterministic waiting time bias signal. This population might represent an intermediate stage of the information flow from the mPFC deterministic waiting time bias neurons to M2 actual waiting time neurons, but we do not know whether the final action timing neurons and the deterministic bias neurons form truly separate populations.

The causal involvement of M2 in waiting decisions was supported by inactivation experiments. M2 inactivation significantly changed the waiting time distribution in all rats, but the direction of the effect was not consistent across rats. This might be because there are both signs of waiting time correlated neurons in M2, and also because there are different types of waiting time coding neurons, namely ramp-to-threshold type and transient correlation type (Figure 3)(Murakami et al., 2014). The possible difference in the composition of different types of neurons affected by muscimol inactivation across animals might result in the variable pattern of changes in the waiting time distribution. M2 inactivation not only changed the waiting time distribution but also impaired general task performance. Strikingly, M2 inactivation reduced the fraction of trials in which the rat visited the reward port after leaving the waiting port, even though the reward was available. This suggests a more general role of M2 in performing the task, such as selecting proper action sequences, which requires planning what action to perform and when to execute it. M2 neural populations active at different trial phases in the waiting task (Murakami et al., 2014) might support this function.

### Incorporating a waiting time bias signal into an integrate-to-bound model for the waiting decision

We previously proposed that the M2 neurons specify the timing of spontaneous actions through an integration-to-bound circuit, where transient waiting time correlated neurons serve as inputs to the integrator and affect the slope of integrator activity (Murakami et al., 2014; Schurger et al., 2012). Once the integrator reaches a fixed threshold level, action is taken, aborting waiting. Extending this circuit, we propose that the deterministic waiting time bias signal in the mPFC is located upstream of the integrator circuit in the M2 and influences the inputs to the integrator. For the waiting time bias signal to be effective in influencing the actual waiting time, it is required for the waiting time bias signal to influence the neurons with positive and negative waiting time correlation in the opposite direction. Although a direct test for this hypothesis is necessary, it is consistent with the noise correlation structure among waiting time correlated M2 neurons found in the previous study (Murakami et al., 2014).

### Behavioral timescale and neural timescale in the prefrontal-motor cortical hierarchy

We found that mPFC and M2 differed in the temporal scale of their waiting time coding. The mPFC encoded only the slowly fluctuating deterministic bias component, but not the rapidly fluctuating stochastic component, while the M2 carried both components. Furthermore, the slowly fluctuating bias signal differed in its temporal dynamics in the two areas: in the mPFC neurons it was a sustained signal, while in M2 neurons it was more transient. These results suggest that the mPFC, compared to the M2, is more specialized in encoding signals that are maintained for a longer time. The results are consistent with previous studies showing that mPFC encodes behavioral variables that have to be maintained for a long timescale across trials, such as a task rule, belief in an environmental model or values of choice options (Bissonette and Roesch, 2015; Karlsson et al., 2012; Rich and Shapiro, 2009; Sul et al., 2010).

In the sensory cortices, it has been shown that the different areas along the cortical hierarchy perform different timescales of information processing (Hasson et al., 2008; Lerner et al., 2011). Different timescales of information processing might be supported by different intrinsic timescales of neural activity across the cortical hierarchy (Murray et al., 2014). Inspired by this idea, we estimated the relative intrinsic timescales of mPFC and M2 dynamics from the decay time constant of the spike count autocorrelation. We found that the mPFC has a longer intrinsic timescale than M2 (514 vs. 373 ms). These findings extend the idea that different timescales of events are handled by cortical areas at different hierarchical levels and suggest that it can be applied to the rodent prefrontal-motor cortical hierarchy: The prefrontal cortex, with its long intrinsic timescale of neural dynamics, carries a signal that needs to be maintained for a long timescale across trials. The motor cortex uses this signal to actually implement an action, which can vary trial-to-trial and moment-to-moment.

In the current study, events from trial history maintained across multiple trials comprise the deterministic waiting bias. Thus it is hard to determine whether the mPFC encodes the deterministic bias in general or it does so only when the deterministic bias has a long timescale. With a sensory cue that signals a probable Tone 2 delay, for example, the deterministic bias could be expected to vary from trial to trial. Using such a cue, we may be able to determine whether the sensory cue biases waiting time through the mPFC or through circuits independent of mPFC. This could help to determine generality/specificity of the deterministic bias encoding in the mPFC.

### Neural circuit for generating stochastic behavioral variability

Decision-making behavior often has a seemingly stochastic component. Even a simple “free-choice” paradigm with a probabilistic reward schedule in highly controlled laboratory settings is often modeled as a probabilistic choice on top of a history-dependent deterministic component (Corrado et al., 2005; Lau and Glimcher, 2005). Our task can be considered a “free temporal choice” paradigm in which a subject is free to abort waiting at any time. In this setting, animals’ behavior can also be modeled as stochastic process on top of a history-dependent deterministic component. The contribution of the deterministic bias component was relatively small (11% of variance explained) and we do not exclude a possibility that other variables that were not included in the model might significantly contribute to determining the animals’ action timing. Yet, importantly, this model was sufficient to reveal a dissociation of encoding schemes in mPFC and M2: mPFC activity encoded the deterministic waiting time bias, but not the actual waiting times. The actual waiting time signal only appears at the level of M2. This suggests that stochastic variability is added somewhere downstream of mPFC and before or at the level of M2. Given the existence of the direct connection from the mPFC to the M2, these results support a two stage model of action timing decisions: mPFC provides a deterministic choice bias signal, which is translated into an actual discrete choice signal by injecting a stochastic trial-to-trial variability at a downstream circuit including M2.

Although stochastic variability in the waiting time is strongly encoded in M2, it remains to be determined whether the variability is generated within M2 itself or it is inherited from other structures. In the songbird, it has been proposed that a frontal cortical structure, the lateral magnocellular nucleus of the anterior nidopallium (LMAN), plays a key role in generating song variability, critical for learning to imitate a tutor song (Fee and Goldberg, 2011). Yet, LMAN is part of a basal ganglia-thalamocortical loop and the importance of thalamic (Goldberg and Fee, 2011) and basal ganglia (Woolley et al., 2014) structures in the generation of song variability has also been suggested. In a rat duration categorization task, trial-to-trial variability in duration judgment is reflected in neural activity within the striatum (Gouvêa et al., 2015). Basal ganglia activity might contribute to variability in waiting times because the waiting decision in our task are likely to be affected by duration judgment. These results raise the possibility that the neural correlates of stochastic variability in M2 might be inherited from other structures in the basal ganglia-thalamocortical loop. Future studies will be necessary to pinpoint the source of stochastic behavioral variability.

## EXPERIMENTAL PROCEDURES

### Animal subjects

All procedures involving animals were either carried out in accordance with National Institute of Health standards and approved by Cold Spring Harbor Laboratory Institutional Animal Care and Use Committee or in accordance with European Union Directive 86/609/EEC and approved by Direcção-Geral de Veterinária. Experiments were performed on 33 male adult Long-Evans hooded rats. These rats are a subset of 37 rats used in our previous study (Murakami et al., 2014). Four rats that performed interleaved blocks of nose-poke waiting trials and lever-press waiting trials were excluded from the analyses, because their data were not appropriate for the waiting time bias fluctuation analysis. Rats had free access to food but water was restricted to the behavioral session and 20 – 30 additional minutes per day.

### Behavioral task

Rats were trained and tested on the waiting task (Figure 1A)(Murakami et al., 2014). The behavioral box contained a wall with three ports (Island motion corporation, Tappan, NY). The waiting port was located at the center and the reward port was located at the side. The side of the reward port was chosen randomly for each rat. The third port was inactive. Entry to and exit from the ports were detected based on an infrared photo-beam located inside each port.

Rats initiated a trial by poking their snout into the waiting port. The first tone, Tone 1 (6 kHz or 14 kHz tone, 80ms) was played, if the rat kept its snout inside the waiting port for T1 delay (= 0.4 s). Tone 1 signaled the availability of a small amount of water reward at the reward port. If the rat moved out before Tone 1, no rewards were available in that trial (short poke trial). If the rat moved out of the waiting port after Tone 1 and visited the reward port, a small water reward (10μl) was delivered through a tube in that port after a 0.5 s delay (impatient trial). If the rat did not respond to Tone 1 and waited with its snout in the waiting port, a second tone, Tone 2 (14k Hz or 6k Hz, differing from Tone 1, 80ms) was played after a certain delay (T2 delay). If the rat visited the reward port after Tone 2, a large reward (40 μl) was delivered after a 0.5 s delay (patient trial). For two of the rats, the small reward was 14 μl and the large reward was 30 μl. There were no signals to the rat that it had exited the waiting port. Re-entrance to the waiting port (multi-poke) was signaled by brief noise burst (60ms) to discourage this behavior. The rat had to visit the reward port within 2s after the initial poke-out in order to collect rewards

T2 delay was drawn randomly from an exponential distribution, whose minimum value was 0.7s and whose mean value was adjusted according to the performance of the animal so that rats succeeded in waiting on about one third of trials. The mean value of the exponential distribution was adjusted every trial; after each short-poke trial or impatient trial, the mean was decreased by 20 ms. After each patient trial, the mean was increased by 40 ms. For the muscimol inactivation experiments, this adjustment procedure was not used.

An inter-trial interval (ITI) period started after the delivery of the reward. During the ITI period, a white noise was played. The time from the initial poke into the waiting port to the end of the ITI was constant, so that the rat could not profit from leaving the waiting port early to start the next trial early. Thus, the optimal strategy to obtain maximal reward in this task was always to wait for Tone 2.

Waiting time was defined as a time from the entry into the waiting port to the movement out of the waiting port (Figure 1A). Movement time was defined as the time from leaving the waiting port to entering the reward port (Figure 1A).

In order to estimate a deterministic waiting time bias of each trial, we used a Cox proportional hazards regression (*coxphfit* in MATLAB statistical toolbox) with waiting times and rewards of past *N* trials as predictors (covariates). Briefly, we estimated the coefficients for the Cox proportional hazards model described as follows:

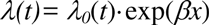

where *λ*(*t*) represents a hazard function (hazard rate of leaving a waiting port), *λ_0_*(*t*) represents a baseline hazard function, that is a hazard function when all the covariates are 0, *β* is a row vector with 2· *N* elements representing Cox coefficients for each covariate and *x* is a 2· *N* element column vector representing covariates, waiting times and rewards of past N trials. Trials from multiple sessions were combined together to estimate 2· *N* coefficients for each rat. For a proportion of explained variance in the Cox regression model, we used a Schemper’s V (Schemper and Henderson, 2000), which measures a variance explained in terms of a survival function.

### Muscimol Inactivation

For the muscimol inactivation experiments, each rat was implanted with 33 gauge stainless steel guide cannulae and dummy cannulae (Plastics One, Roanoke, VA)(Felsen and Mainen, 2008; Narayanan et al., 2006) targeted to M2 (4.2 mm anterior to and 1.75 mm lateral to and 2.0 mm deep from Bregma) or mPFC (3.2 mm anterior to, 0.75 mm lateral to and 3.6 mm deep from Bregma) bilaterally (Paxinos and Watson, 2005). The guide cannulae tips were located 0.5 mm dorsal to the target depth, so that the internal cannulae extending 0.5 mm beyond the guide cannulae tips would be placed at the target sites during injections. Rats were allowed to recover for 5 days before water restriction resumed and the inactivation sessions began. Prior to each session, the rat was anesthetized with 2% isoflurane. Hamilton syringes (Hamilton Company, Reno NV), connected with 33 gauge internal cannulae (Plastics One) through plastic tubes and mounted on an infusion pump (Harvard Apparatus, Holliston, MA), were used to administer 0.5 μl of either muscimol or saline at a rate of 0.25 μl/min bilaterally. Animals recovered for 1 hour before beginning the 1 hour behavioral session. For mPFC inactivation, rats were injected with 0.2 μg/μl muscimol. For M2 inactivation, rats were injected with 0.05 μg/μl muscimol.

### Neural recording

For the recording experiments, each rat was implanted with a drive (Island motion corporation, Tappan, NY) containing 10 to 24 movable tetrodes targeted to the mPFC (3.2 mm anterior to and 0.75 mm lateral to Bregma) or M2 (3.2 – 4.7 mm anterior to and 1.5 – 2.0 mm lateral to Bregma) contralateral to the side of the reward port (Paxinos and Watson, 2005). For mPFC targeting, tetrodes were tilted by 12 degrees. Rats were allowed to recover for 5 days before water restriction resumed and the recording sessions began.

Individual tetrodes consisted of four twisted polyimide-coated nichrome wires (H.P. Reid, Inc., Palm Coast, FL; single-wire diameter 12.5 μm) gold-plated to 0.2 – 0.5 MΩ impedance at 1 kHz. The tetrodes were coated with DiI (Molecular Probes, Eugene, OR) to visualize the tetrode tracks in a histological examination. Electrical signals were amplified and recorded using the NSpike data acquisition system (L. M. Frank, University of California, San Francisco, and J. MacArthur, Harvard University Electronic Instrument Design Lab). Multiple single units were isolated offline by manually clustering spike features derived from the waveforms of recorded putative units using MCLUST software (A.D. Redish). Tetrode depths were adjusted before or after each recording session in order to sample an independent population of neurons across sessions. The locations of tetrode tips during each recording session were estimated based on their depth and histological examination based on electrolytic lesions and the visible tetrode tracks. All single-units recorded from mPFC (including the anterior cingulate cortex, prelimbic cortex and infralimbic cortex) or M2 were included in the analysis. Rats performed 1 session per day, and a total of 306 neurons were recorded from M2 over 61 recording sessions from 5 rats and 126 neurons were recorded from mPFC over 26 sessions from 3 rats. When we excluded single-units suspected to be recorded during more than one session, as judged from the spike waveform and the firing pattern, the results remained qualitatively the same (data not shown). Electrolytic lesions were produced after the final recording session (15 μA of cathodal current, 10 s).

### Histology

In order to verify the location of the injection cannulae or the tetrodes, rats were then deeply anesthetized with pentobarbital and perfused transcardially with 4% paraformaldehyde. The brain was sectioned at 50 μm. Slices were stained with Cresyl violet solution. For the neural recording rats, every other slices were prepared for fluorescent observation to examine the fluorescent tracks made by DiI-coated tetrodes.

### Inactivation data analysis

For the muscimol inactivation experiments in Figure 2, we measured, for each rat and each session, the median waiting time, fraction of patient trials, median movement times of impatient or patient trials, fraction of reward port visit trials of impatient or patient trials. On population rat data, we performed repeated measures ANOVA across 3 sessions, followed by t-tests comparing saline-pre (Sal-pre) vs. muscimol (Mus) and muscimol (Mus) vs. saline-post (Sal-post) sessions. We defined a significant muscimol effect, here and elsewhere, as significant differences (*P* < 0.05) in both comparisons in the consistent direction. Thus, *P*-value was not corrected for multiple comparisons. Parametric tests were used due to the relatively small sample sizes.

In addition to population statistics, we also performed statistical tests on individual rats (black or white circles in Figure 2). As for the fraction of patient trials and fraction of reward port visit trials, we performed chi-squared test on contingency tables on data across 3 sessions, followed by Marascuillo procedures comparing Sal-pre vs. Mus and Mus vs.Sal-post sessions. As for the waiting times and movement times, we performed one-way ANOVA on data across 3 sessions, followed by two-sample t-tests comparing Sal-pre vs. Mus and Mus vs. Sal-post sessions. To test differences in the waiting time distributions across sessions, we also performed Kolmogorov-Smirnov tests comparing Sal-pre vs Mus and Mus vs. Sal-post sessions.

### Neural data analysis

All data analysis was performed with custom-written software using MATLAB (Mathworks, Natick, MA). No statistical methods were used to pre-determine sample sizes. But our sample sizes were similar to those reported in previous studies. Two-sided tests were used for all the statistical tests unless stated otherwise. All the spike density functions (SDFs) were generated with a Gaussian filter (s.d. = 50 ms) only for visualization purpose, unless stated otherwise.

### Waiting time correlation

In order to search for waiting time correlated activity in Figure 3, we performed the same analyses as in our previous study (Murakami et al., 2014). For these analyses, only the impatient trials were used because in the patient trials, the response was triggered by the tone. Because in the multi-poke trials, we did not know whether the rat intended to leave the waiting port or just failed to wait unintentionally, we excluded this type of trials from the neural analysis. We also excluded the trials in which the rat did not visit the reward port within 2s after the initial poke-out.

For the ramp-to-threshold analysis, only those neurons which showed different firing rates during the delay period (from 0.4 s after poke-in to 0.4 s before poke-out) and the poke-out period (last 0.4 s before poke-out) were analyzed (Wilcoxon signed-rank text, *P* < 0.01, “ramp-up” or “ramp-down” neurons). Neurons with less than 10 long waiting trials (more than 1.2 s waiting time) were pre-excluded before the selection process because the delay period firing rate could not be estimated reliably. Impatient trials were divided into 10 groups based on the waiting time, with equal (or different by 1) number of trials per group. Spike trains were smoothed with a causal filter (EPSP-like filter, which is a multiple of 2 exponentials, one rising exponential with τ = 1 ms, and the other falling exponential with τ = 40 ms) to generate a spike density function (SDF) for each group. The threshold crossing time was determined from the SDFs. While ramp-up neurons were tested with threshold crossing with positive slope, ramp-down neurons were tested with negative slope. Threshold crossing time was detected when the SDF first crossed the threshold and stayed above (or below for ramp-down neuron) the threshold for more than 20 ms. The threshold crossing time was defined as the end point of this 20 ms. The search for threshold crossing was started at the trough time between 0 and 0.4 s from the poke-in (peak time for the ramp-down neurons) of the average of the 10 SDFs and ended at 0.4 s after the waiting time of the longest waiting time trial. For each neuron, we tested 10 different firing rate thresholds, determined as follows: The lowest threshold was set at the lowest firing rate at which threshold crossing occurred in at least 9 out of 10 SDFs. The highest threshold was set at the highest firing rate at which threshold-crossing occurred at least in 9 SDFs. Eight intermediate thresholds were equally-spaced in between the lowest and the highest thresholds. At each threshold level, we calculated correlation between the mean waiting times against the threshold crossing times. A ramp-to-threshold type predictive neurons was defined as a neuron which showed, 1) significant correlation coefficient between the threshold crossing time and the waiting time in at least 2 of the 10 thresholds (corrected for multiple comparisons), 2) positive prediction time (defined below) in at least 2 significant thresholds and 3) the regression slope for threshold crossing time against the waiting time close to unity (value between 0.8 and 1.2) in at least 1 significant threshold with positive prediction time. Prediction time was defined for each threshold as an average of [waiting time – time to cross threshold] across different waiting time groups. In order to estimate the significance of fraction of ramp-to-threshold type neurons, we ran a permutation test by randomly permuting the waiting times and neural data across impatient trials. We repeated this procedure 1000 times to estimate the probability of obtaining the observed fraction of significant neurons by chance.

We also ran a version of ramp-to-threshold analysis with less strict criteria. In this analysis, the step of pre-selecting neurons with different firing rates between the delay period and the poke-out period was omitted. Each neuron was tested with threshold crossing with both positive and negative slope. A ramp-to-threshold type neuron was defined as a neuron which showed, in at least 1 of the 10 thresholds, significant correlation coefficient between the threshold crossing time and the waiting (corrected for multiple comparisons) with positive prediction time.

For the transient correlation analysis, we excluded ramp-to-threshold type neurons, because in this analysis, we looked for a different class of waiting-time predictive activity from the ramp-to-threshold type neurons. The correlation coefficient between the firing rate and the waiting time was examined for non-overlapping 0.4 s time window, starting from 1.2 s before the waiting port entry. For each time window, the trials in which the rat had already exited the waiting port by the end of the analysis time window or within 0.2 s after the end of the time window were excluded from the analysis, in order to exclude potential movement correlates. Time windows with less than 10 trials after the exclusion were not analyzed, because the correlation could not be estimated reliably. Because the significance was tested in multiple time windows, we corrected the significance level using Bonferroni correction. In order to estimate the significance of fraction of firing rate correlated neurons, we ran a permutation test by shuffling the impatient trials 1000 times.

### Waiting time bias correlation

In Figure 4, we performed a waiting time bias correlation analysis in the same way as we did for the transient correlation analysis, but the waiting time bias, instead of the actual waiting time, was used. The waiting time bias was estimated using the Cox regression analyses. The waiting time and reward history up to 5 trial back were included in the Cox regression model, but including up to 3 or 10 trial back did not change the results qualitatively (data not shown). We used only the impatient trials so that the fraction of significant neurons was comparable to the actual waiting time correlation analysis and the multiple regression analysis below. In order to estimate the significance of fraction of firing correlated neurons, we ran a permutation test by shuffling the impatient trials 1000 times. Additionally, in order to maintain slow fluctuations of neural activity and waiting times in shuffle data but to destroy the relationship between those, we also performed session shuffling. In order to control for the number of trials used for the shuffling analysis and the original analysis, we only used a fixed number of trials (from trial 1 to trial 200). Neurons with fewer than 200 trials were excluded from this analysis.

In Figure 5, we performed multiple linear regression analyses, with waiting time biases and residual waiting times as independent variables to predict neuron’s firing rates of a 0.4 s bin. Only impatient trials were used for the analysis. The residual waiting times were calculated by subtracting the waiting time bias from the actual waiting time. The waiting time bias, residual waiting time bias and firing rate were z-score normalized before performing the multiple linear regression. *P*-value was calculated for each predictor at each time bin. Neurons significantly correlated with waiting time biases or residual waiting times were defined as a neuron that had, at least in one of the analyzed time bins, a regression coefficient significantly different from 0 (*P* < 0.05, after correcting for multiple comparisons for multiple time bins).

### Demixed principal component analysis

In Figure 3G and 3H, we performed demixed principal component analysis (dPCA) as described previously (Kobak et al., 2016). The objective of dPCA here is to project activity of a neural population into reduced dimensions that capture the variance in activity due to difference in waiting times across trials, but not due to time-varying activity within a trial. Briefly, for each neuron, we created 6 poke-in aligned SDFs of impatient trials with 6 different waiting time groups. The group separation boundary started from 0.4 s, with 0.2 s increment, and ended at 1.6 s. Neurons without at least two impatient trials for each waiting time group were excluded from the analysis. The remaining 278 M2 neurons and 114 mPFC neurons were used for this analysis. To create SDFs, activity from −1.2 s before the poke-in till the shortest waiting time of each group (for example, 0.4 s for the first group) was convolved with Gaussian kernel with 50 ms standard deviation, averaged across trials and sampled at 10 ms resolution. For each neuron, first, we concatenated the SDFs from different groups and z-score normalized. The data matrix, X, was organized so that each row containss the normalized concatenated SDFs for each neuron and different rows represent different neurons. The data matrix, X, was decomposed into the time-varying component X_*t*_ and the remaining waiting time comoponent X_*wt*_, so that X = X_*t*_ + X_*wt*_. X_*t*_ was generated by averaging, for each neuron, SDFs across waiting time groups at each time point. (Note that in later times, there are fewer groups to be averaged. Also note that X_*t*_ has the same dimension as X, with the same values repeated across groups). The remaining waiting time component X_*wt*_ was given by X_*wt*_ = X − X_*t*_.

Lastly, we performed demixed PCA by minimizing the loss function, L, given below:

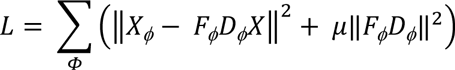

with constraints of orthonormality of F_ϕ_ (but not necessarily for D_Φ_) and rank(F_Φ_D_Φ_ = N (we used N = 10). Here, the summation is over different components Φ = t, wt, ‖·‖ indicates Frobenius norm, F_Φ_ and D_Φ_ are the encoder and decoder matrices, respectively, and μ is a regularization parameter. For given μ, the minimization can be performed separately for each Φ, which amounts to following steps:

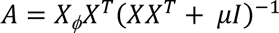

where X^T^ is a transpose of X, I is the identity matrix. We obtain F_Φ_ = U and D_Φ_ = U^T^A, where U is the first N principal axis of AX. Regularization parameter was determined by 10 times cross-validation (see Kobak et al., 2016 for details). Presented data are projections of X with the first few rows of the decoder matrix D_wt_.

Different numbers of neurons are used in dPCA for M2 (278 neurons) and mPFC (114 neurons). In order to control for this difference, we subsampled the 114 M2 neurons without replacement and performed the dPCA (Figure S2C and S2D). We iterated this procedure 1000 times and evaluated the strength of wait time coding based on correlation coefficient between the dPCA socres and waiting time. The strong and ordered wait time signal was still observed with subsampled M2 population.

In Figure 4, the dPCA targeting the waiting time bias signals instead of the waiting time signals was performed. The impatient trials were separated into 6 groups according to the waiting time bias. The group separation boundaries started from 2.5 percentile of the waiting time bias of the session and ended at 97.5 percentile of the waiting time bias. Five intermediate boundaries were chosen to equally space the time between the start and end values. To create SDFs, activity from −1.2 s before the poke-in till 0.4 s after the poke-in was used. The waiting time bias component was demixed from the time-varying component in the same way as described above.

### Intrinsic timescale

We measured the intrinsic timescale of each brain area as reported previously (Murray et al., 2014). We used neural activity from 2 s time window before the poke-in, binned into twenty 100 ms bins. Across-trial spike-count correlation was calculated for each pair of bins, yielding 20 × 19 correlation coefficients for each neuron, excluding time lag = 0. The data were pooled across neurons from the same brain area. The decay of autocorrelation was fit by an exponential function with an offset,

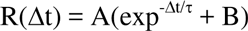

where R(Δt) is an autocorrelation value at time lag, Δt, A (Scaling factor), B (offset), τ (decay constant) are the free parameters. For fitting, we used *nlinfit* function in MATLAB (Mathworks, Natick, MA) with initial values of A = 1, B = 1, τ = 1000 ms. Only the decaying part of the autocorrelation function was used for fitting. Thus, autocorrelation at time lag = 100ms was excluded for fitting the mPFC data.

Confidence intervals were obtained with bootstrapping, by resampling neurons with replacement 1000 times. Statistical difference between brain areas is estimated with permutation test, by randomly shuffling neurons between brain areas 2000 times.

We also calculated spike-count correlation between two consecutive trials, using a single 2 s time window before the poke-in. Confidence intervals and statistical difference were estimated in the same way as described above.

### Coherency analysis

For the coherency analysis in Figure 6, we used, for each neuron, 4 time bins, from −1.2 s till 0.4 s from poke-in. For each time bin, we have actual waiting times and the firing rate across trials. We z-score normalized each data. We then simply calculated a coherency between these two signals with a multi-taper method using a chronux package (http://chronux.org). Time-bandwidth product 16 and 31 tapers were used. We used all types of trials (short-poke, impatient and patient trials) for this analysis. Trials in which a period after poke-out overlapped with the analysis time bin were excluded. For each neuron, we only used time bins until 0.4 s from poke-in, because time bins after this would have many excluded trials and coherency might not be estimated accurately. Absolute coherency was averaged across time bins and across all the neurons from each brain area. Session shuffling was done in the same way as described above.

### Waiting time bias correlation across trial phases

For the across trial phase analysis in Figure 7, trial phases were defined as follows: Pre-wait phase was −1.2 – 0 s from the poke-in. Wait-start phase was 0 – 0.4 s from the poke-in. Wait-end phase was −0.4 – 0 s from the poke-out. Move phase was from the poke-in till the poke-out. Water-delay phase was from the reward-poke-in till the water delivery (0.5 s from the reward poke in). Water/ITI phase was 0 – 2 s from the water delivery. The correlation between waiting time bias and the firing rate was calculated for each phase. Neurons were pre-selected if they exhibited significant correlation at the wait-start period (*P* < 0.05, not corrected for multiple comparisons). If the correlation at the wait-start phase was negative, the sign of correlations for all the trial phases for this neuron were flipped, before across-neuron averaging. We also performed the same analysis, but selected neurons according to the waiting time bias correlation at different trial phases. The results were qualitatively the same (Figure S3).

## AUTHOR CONTRIBUTIONS

M.M. and Z.F.M designed the experiments and M.M. performed the experiments. M.M., H.S. performed the analyses and M.M., H.S. Y.L. and Z.F.M. discussed the analyses. M.M. and Z.F.M. wrote the manuscript

## ACKNOWLEDGEMENTS

We thank Maria Inês Vicente, Gil M. Costa, Mark Terrelonge for help in behavioral training and electrophysiology, Nicole Horst and Mark Laubach for advice on muscimol inactivation procedures, Dmitry Kobak and Pietro Vertechi for advice on demixed principal component analysis, Barry Burbach and Mauricia Vinhas for technical assistance, Fanny Cazettes, Luca Mazzucato, Gautam Agarwal, Eran Lottem, Cindy Poo, Bassam Atallah for helpful comments on the manuscript. This work was supported by European Research Council (250334 and 671251, Z.F.M), Simons Foundation (325057, Z.F.M), Champalimaud Foundation (Z.F.M), the Israel Science Foundation (757/16, Y.L.), the Gatsby Charitable Foundation (Y.L.), Fundaçao para a Ciência e a Tecnologia (SFRH/BPD/46314/2008, M.M.), the Uehara Memorial Foundation (M.M.) and Fundaçao Bial (127/08, M.M.).

## FIGURE LEGENDS

**Figure S1.**
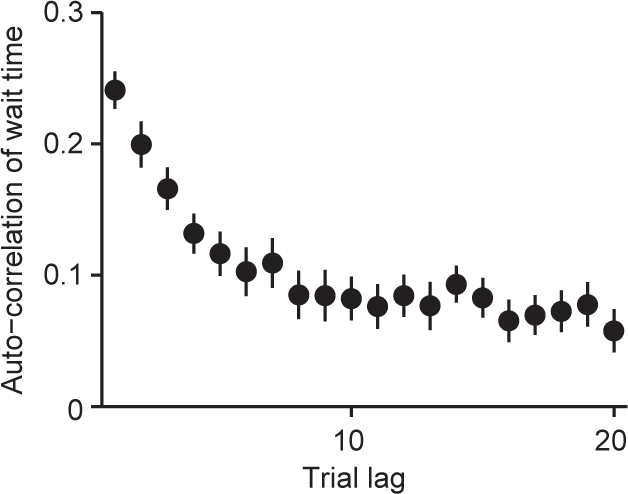
Auto-correlation of Waiting Time across Trials, Related to Figure 1. Auto-correlation of waiting time across trials was calculated for each rat combined across sessions. Shown is a mean and S.E.M. across rats (*N* = 33 rats).

**Figure S2.**
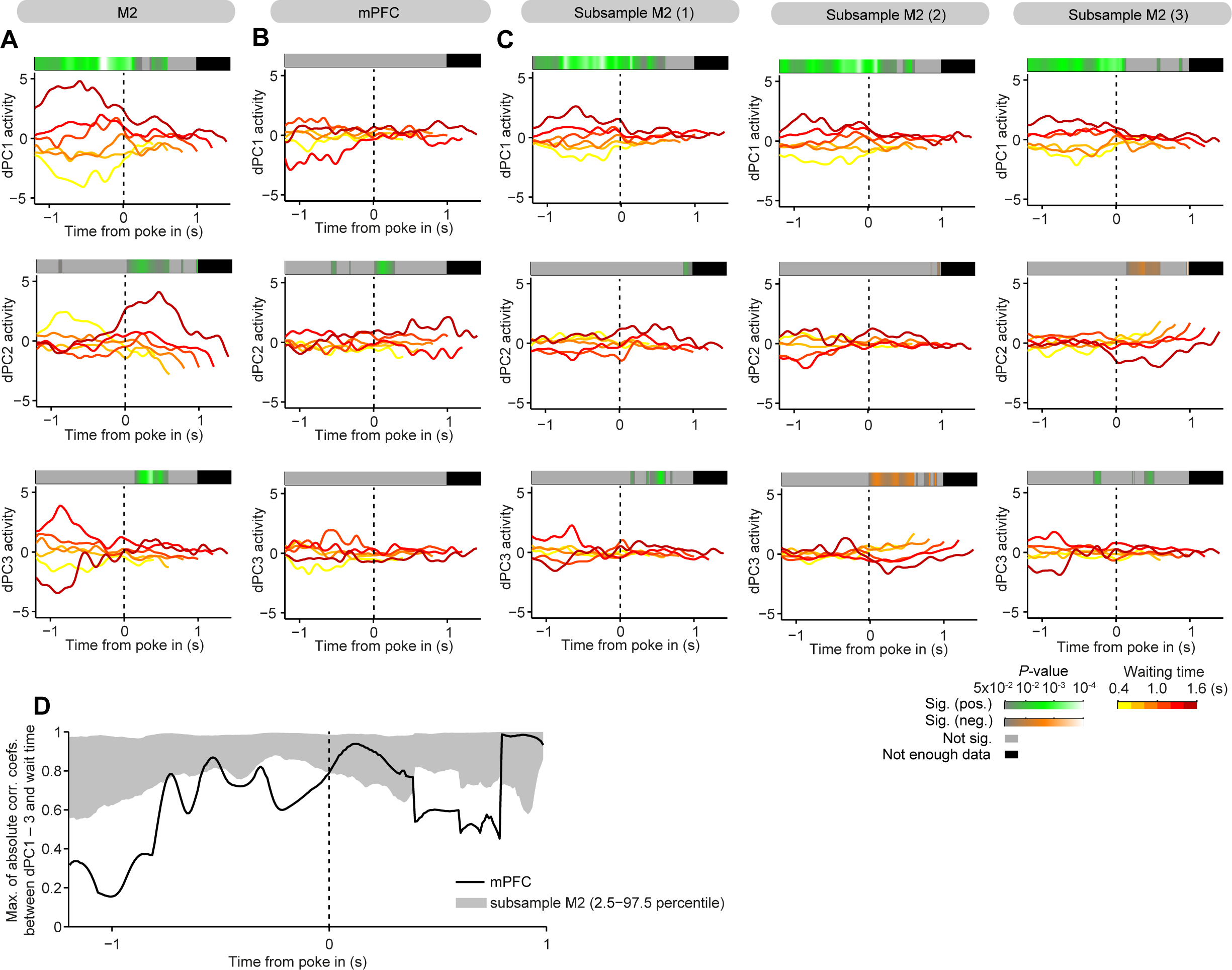
More Components of Demixed Principal Component Analyses and Neuron Number Control Analysis, Related to Figure 3. (A) The same as Figure 3G, but the activity of dPC1 – 3 are shown from top to bottom. (B) The same as (A), but for mPFC neurons. (C) The same as (A), but for subsampled M2 population. Three examples of random subsampling were shown from left to right. (D) Maximum of absolute correlation coefficients between dPC1 – 3 activities and the waiting time is plotted as a function of time relative to the poke-in. The solid line represents mPFC data and the gray area represents 95 percentile range (2.5 – 97.5 percentile) of 1000 iterations of subsampled M2 data.

**Figure S3.**
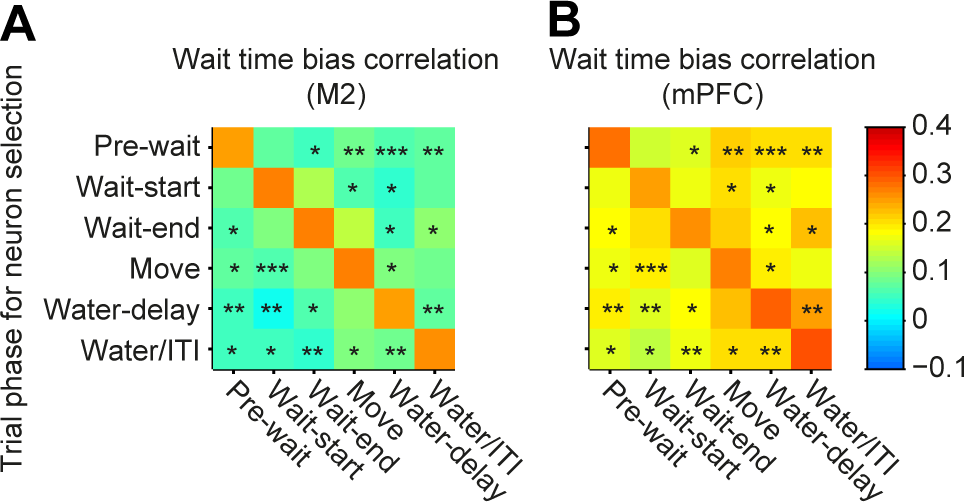
Temporal Dynamics of the Waiting Time Bias Signal in M2 and mPFC Does Not Depend on the Neuron Selection Criteria, Related to Figure 7. (A) The same analysis as Figure 7D, but using different phases for selecting the waiting time bias correlated neurons. For each row, neurons are selected based on the waiting time bias correlation at a given trial phase labeled on the y-axis (Figure 7D corresponds to the second row). Shown are mean correlation coefficients across neurons (*N* = 49, 40, 46, 58, 37, 66 M2 neurons from top to bottom). Asterisks indicate significant difference between M2 and mPFC (*: *P* < 0.05, **: *P* < 0.01, ***: *P* < 0.001). (B) The same as (A) but for mPFC neurons (*N* = 25, 25, 22, 26, 20, 25 mPFC neurons from top to bottom).

